# Intracellular signaling by the *comRS* system in *Streptococcus mutans* genetic competence

**DOI:** 10.1101/291088

**Authors:** Simon A.M. Underhill, Robert C. Shields, Justin R. Kaspar, Momin Haider, Robert A. Burne, Stephen J. Hagen

## Abstract

Entry into genetic competence in streptococci is controlled by ComX, an alternative sigma factor for genes that enable the import of exogenous DNA. In *Streptococcus mutans* the immediate activator of *comX* is the ComRS quorum system. ComS is the precursor of XIP, a seven-residue peptide that is imported into the cell and interacts with the cytosolic receptor ComR to form a transcriptional activator for both *comX* and *comS*. Although intercellular quorum signaling by ComRS has been demonstrated, observations of bimodal expression of *comX* suggest that *comRS* may also function as an intracellular feedback loop, activating *comX* without export or detection of extracellular XIP. Here we use microfluidic and single cell methods to test whether ComRS induction of *comX* requires extracellular XIP or ComS. We find that individual *comS*-overexpressing cells activate their own *comX*, independent of the rate at which their growth medium is replaced. However, in the absence of lysis they do not activate *comS*-deficient mutants growing in co-culture. We also find that induction of *comR* and *comS* genes introduced into *E. coli* cells leads to activation of a *comX* reporter. Therefore ComRS control of *comX* does not require either the import or extracellular accumulation of ComS or XIP, or specific processing of ComS to XIP. We also find that endogenously and exogenously produced ComS/XIP have inequivalent effects on *comX* activation. These data are fully consistent with intracellular positive feedback in *comS* transcription as the origin of bimodal *comX* expression in *S. mutans*.

**Importance:** The ComRS system can function as a quorum sensing trigger for genetic competence in *S. mutans.* The signal peptide XIP, which is derived from the precursor ComS, enters the cell and interacts with the Rgg-type cytosolic receptor ComR to activate *comX*, which encodes the alternative sigma factor for the late competence genes. Previous studies have demonstrated intercellular signaling via ComRS, although release of the ComS or XIP peptide to the extracellular medium appears to require lysis of the producing cells. Here we test the complementary hypothesis that ComRS can drive *comX* through a purely intracellular mechanism that does not depend on extracellular accumulation or import of ComS or XIP. By combining single-cell, coculture and microfluidic approaches we demonstrate that endogenously produced ComS can allow ComRS to activate *comX* without requiring processing, export or import. These data provide insight into intracellular mechanisms that generate noise and heterogeneity in *S.mutans* competence.

## Introduction

*Streptococcus mutans* inhabits human oral biofilms and is a primary etiological agent of dental caries (1). Many of the behaviors that facilitate the growth, competition, stress tolerance, and virulence of *S. mutans* are linked to the regulation of genetic competence, a transient physiological state during which the organism can internalize DNA from its environment (2-7). The competence pathway of *S. mutans* is complex, as it receives input from extracellular peptide signals and environmental cues as well as regulatory feedback (8-14). Consequently, several elements of the mechanism and dynamics of the pathway are not well understood.

*S. mutans* initiates entry into the competent state by increasing the transcription of the *comX* gene (sometimes referred to as *sigX*), which encodes an alternative sigma factor that is required for the expression of approximately 30 late competence genes (15,16). Expression of *comX* can be induced by the peptides CSP (<u>c</u>ompetence <u>s</u>timulating <u>p</u>eptide) and XIP (*sig*<u>X</u>-<u>i</u>nducing <u>p</u>eptide), and the efficacy of these peptides is strongly influenced by environmental conditions. CSP is derived by cleavage of a 21-residue peptide from ComC and exported through an ATP-binding cassette transporter. It is further processed to the active 18-residue peptide by the SepM protease (17). Extracellular CSP is detected by the two-component signal transduction system ComDE, with the phosphorylated response regulator ComE activating genes for bacteriocin synthesis and immunity. ComE does not directly activate *comX*, but affects *comX* indirectly, via a pathway that is not completely understood (18).

The immediate regulator of *comX* in *S. mutans* and in streptococci of the salivarius, bovis and pyogenic groups is the ComRS system. ComR is an Rgg-like cytosolic transcriptional regulator, and the type II ComS of *S. mutans* is a 17-residue peptide (19). The C-terminus of ComS contains the 7-residue small hydrophobic peptide XIP. Extracellular XIP is imported by the Opp permease and interacts with ComR to form a transcriptional activator for both *comX* and *comS* (19, 20). Notably, the *S. mutans* competence pathway contains at least two positive feedback loops, as XIP/ComR activates *comS* and ComX activates *comE* expression (19, 20).

An intriguing property of *S. mutans* competence is that although exogenous CSP and XIP can both activate *comX* and induce transformability, they do so under different environmental conditions and they elicit qualitatively different behaviors in the expression of *comX* (11). Exogenous CSP elicits a bimodal response in which less than half of the population activates *comX*, whereas exogenous XIP elicits a unimodal response in which all cells in the population activate *comX*. Further, CSP activates *comX* only in complex media containing small peptides; this activation requires that cells carry an intact *comS* but it does not require the Opp permease (11). By contrast, exogenous XIP activates *comX* only in defined media lacking small peptides (11, 21); this activation requires that cells carry the *opp* gene but does not require *comS* (19). Therefore, although exogenous XIP can activate *comX* in a strain lacking *comS*, the bimodal *comX* response to CSP requires an intact *comS* gene.

The observation that competence in several streptococcal species is directly stimulated by an extracellular ComS-derived peptide suggests that ComRS constitutes a novel type of Gram-positive quorum signaling system, in which the ComS-derived signal XIP is processed and secreted, accumulates in the extracellular medium, and is then reimported. This interpretation in *S. mutans* is supported by several experimental observations. First, cells that carry *opp* take up exogenous XIP (in defined medium) and activate *comX* with high efficiency (19, 21). Second, exogenous synthetic XIP is dramatically more effective in stimulating transformability than is exogenous full-length ComS (19). Third, filtrates of *S. mutans* cultures grown to OD550 = 0.4 in defined medium were able to stimulate a P*comX* reporter strain (21). Moreover, LC-MS/MS analysis of supernatants of *S. mutans* cultures grown to high density (22, 23) detected μM quantities of XIP. Fourth, a transposon mutagenesis screen in *S. pyogenes* identified the widely conserved *pptAB* ABC transporter as a possible exporter of short hydrophobic peptides of the ComS type (24), raising the possibility that *S. mutans* may also possess dedicated mechanisms for processing and export of ComS/XIP.

However such mechanisms have not yet been identified in *S. mutans*. There is evidence that the Eep membrane protease facilitates the processing of *S. thermophilus* ComS (25), but Eep did not affect processing of ComS of *S. mutans* (22). Further, although *S. mutans* appears to encode a gene product with a fairly high degree of homology to PptAB of *S. pyogenes*, deletion of the apparent *pptAB* genes had only a weak effect on competence induction in mid-exponential phase cultures of *S. mutans* (24). Consequently, the processing of ComS to XIP remains uncharacterized. The import of XIP presents an additional puzzle for ComRS quorum signaling because the permease Opp is required for XIP to activate *comX*, but is not required for activation by CSP.

We recently demonstrated that XIP can function as a diffusible, intercellular signal (26) in *S. mutans*. A ComS producing strain was able to induce *comX* response in co-cultured cells lacking *comS*, in the absence of physical contact. However intercellular signaling was greatly impaired by deletion of the gene *atlA* encoding the primary autolysin. Therefore, lysis and not active transport appears to be the primary route by which *S. mutans* externalizes the diffusible signal.

Here we use a combination of microfluidic and coculture methods to test the complementary hypothesis that ComRS can also activate *comX* by an intracellular mechanism, through endogenous ComS production, and that this mechanism does not specifically require processing of ComS/XIP or its extracellular accumulation.

## Results

### An intact copy of *comS* alters the *comX* response to exogenous XIP

A previous study found that even when a *comS* deletion (*ΔcomS*) and wild type background were supplied with high concentrations of synthetic XIP, the transformation efficiency of the *ΔcomS* strain was only half as great as that of the wild type (19). Fig. 1A-1B shows a similar experiment performed with individual cells carrying a P*comX-gfp* plasmid-borne reporter. *S. mutans* UA159 (wild type) and Δ*comS* genetic backgrounds were imaged while adhered within a microfluidic chamber and supplied with a constant flow of defined medium (FMC) containing synthetic XIP. Although both strains respond to exogenous XIP, the Δ*comS* strain consistently showed roughly 1.5-fold lower P*comX* activity than the wild type, even at saturating XIP concentrations. The threshold for *comX* response occurred at a roughly 2-fold lower XIP concentration in UA159 than in the Δ*comS* strain (Fig. 1B). Therefore, the deletion of *comS* elevated the threshold for a response to extracellular XIP and reduced the overall response at saturation.

**Fig. 1:**
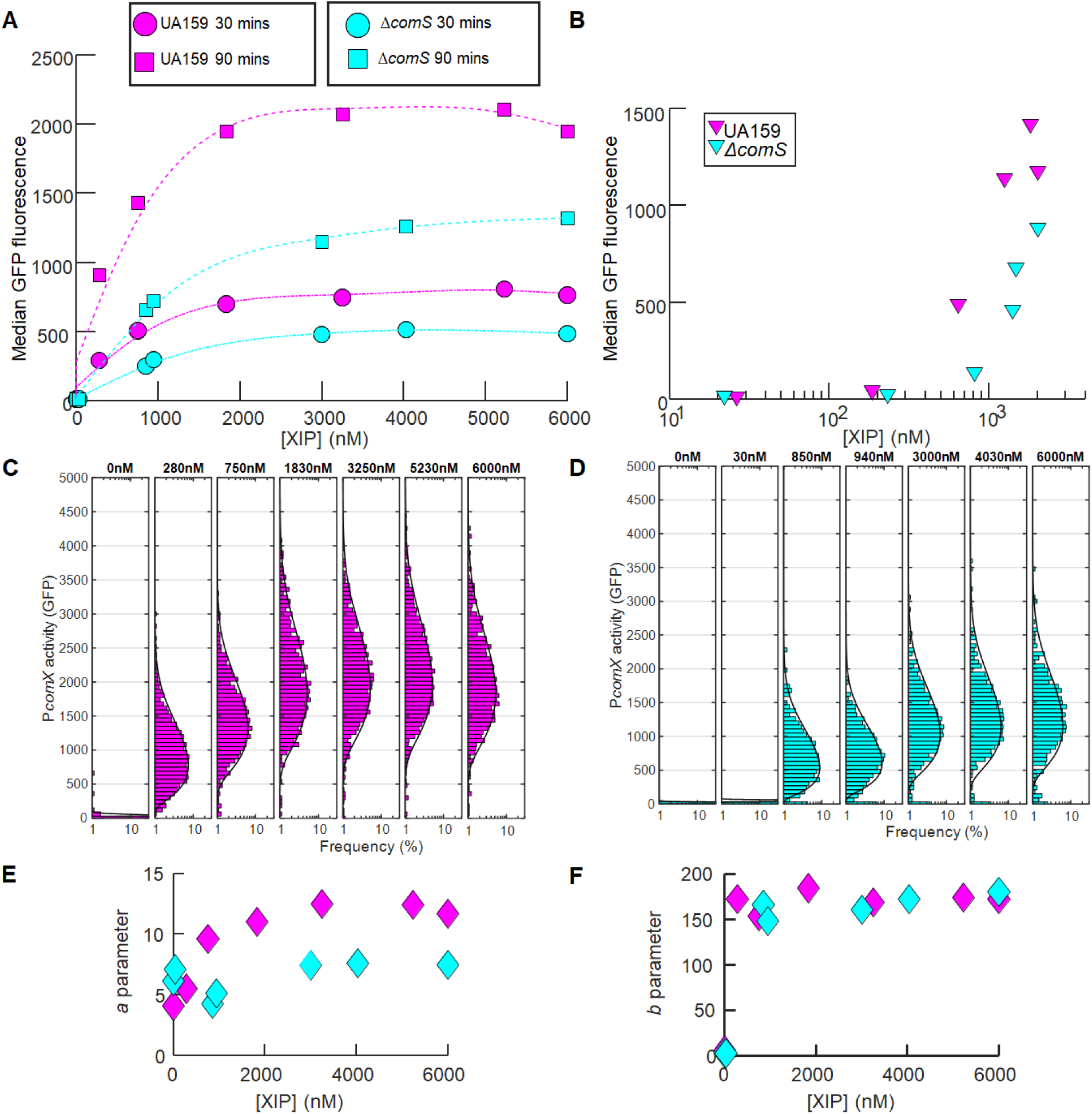
*comS* deletion is not fully complemented by synthetic XIP. (A) Comparison of P*comX-gfp* activity in *S. mutans* cells of the UA159 (wild type) background (magenta) and Δ*comS* mutant background (cyan). The median GFP fluorescence is shown for cells that were supplied with continuous flow of exogenous synthetic XIP in microfluidic chambers. Data are shown at 30 minutes (circles, dash-dot lines) and 90 minutes (squares, dashed lines) of flow. The smooth curves are spline fits to the data. (B) Median GFP levels in the two strains at the 90 minute time point of the flow experiment. Also shown are the histograms of the individual cell P*comX-gfp* reporter activity, versus exogenous XIP concentration, for (C) the UA159 background and (D) the Δ*comS* background. Solid black curves in (C) and (D) show the best fit gamma probability distribution for each histogram. A cutoff of 40 units of P*comX*-*gfp* fluorescence has been applied to exclude background autofluorescence. (E) Parameter *a* of the (two parameter) gamma probability distribution, obtained from fits in (C)-(D), reflecting the ratio of transcription rate to protein degradation rate, and (F) the distribution parameter *b*, reflecting the ratio of translation rate to the rate of mRNA degradation. In (E)-(F) cyan indicates the Δ*comS* mutant, and magenta indicates the UA159 background.

The deletion of *comS* also affected cell-to-cell variability (noise) in *comX* expression. Figs. 1C and 1D show that the histograms of reporter fluorescence differ in UA159 (wild type) and Δ*comS* cells. UA159 showed a generally broader (noisier) *comX* response than did Δ*comS*. We quantified this difference by fitting the histograms to a gamma distribution Γ(*n | a,b*), a two-parameter probability distribution that can be used to model cell-to-cell variability in *n*, the copy number for a bacterial protein (27). In a simple physical model, the parameter *a* of the gamma distribution is related to the number of mRNAs produced during the cell division time, while *b* is related to the number of protein copies produced per mRNA transcript (28). As shown in Figs. 1E and 1F, the UA159 background has a roughly 2-fold higher value for parameter *a* (transcription rate), while parameter *b* (translation) is similar for the two strains. As this difference persists even at XIP concentrations exceeding 1 μM, these data show that deletion of *comS* significantly affects *comX* expression, even when excess extracellular XIP is provided.

### Fluid replacement did not alter induction of *comX* in a *comS* overexpressing strain

To test whether *comX* activation requires cells to import extracellular XIP from their environment, we examined the effect of fluid flow rate in cells that overexpressed *comS* from the strong P23 promoter on the plasmid pIB184 (26, 29) while adhered in a microfluidic flow chamber. Cells carrying the 184*comS* overexpression plasmid produce significantly higher levels of *comS* mRNA (Fig. S1) and activate *comX* in defined medium lacking exogenous CSP or XIP (26). We anticipated that if cells were immobilized and supplied with a continuous flow of fresh medium, high flow rates would remove XIP (or ComS) that was released, leading to diminished *comX* activity. In order to prevent XIP import via Opp, cells were supplied with a flow of complex medium (BHI) lacking XIP or CSP. We loaded 184*comS* P*comX-rfp* cells, which carry both the *comS* overexpression plasmid and a plasmid-borne P*comX*-*rfp* reporter, into four different microfluidic flow chambers. Chambers were supplied with fresh complex medium flowing at rates between 0.02 ml h^-1^ and 1 ml h^-1^. These flow rates were sufficient to completely replace the growth medium within each chamber on time intervals ranging from 6 seconds to 10 minutes. We also studied (I) a P*comX-rfp* reporter in a UA159 background (negative control) and (II) a ComS*-*overproducing strain lacking a start codon (ATG point-mutated to AAG) on its chromosomal *comS* (184*comS* P*comX-rfp* Δ*comS*).

Figs. 2A shows that P*comX* was not activated in the UA159 background (leftmost column). In contrast, the strain overexpressing *comS* and harboring an intact chromosomal copy of *comS* showed a highly heterogeneous response, indicating that a subpopulation of these cells strongly activated *comX* in the flowing complex medium. Further, the rate of fluid flow had no effect on their *comX* expression. Fig. 2B shows the fluorescence of the individual cells for which the signal exceeded the maximum P*comX* activity (roughly 100 fluorescence units) seen in the UA159 negative control. Rather than declining at high flow rates where the medium was rapidly replaced, the median *comX* activity in the ComS*-*overproducing cells actually showed a very slight increase, smaller than the cell-to-cell variability. These data show that overexpression of *comS* can allow *S. mutans* to activate *comX*, even in complex medium which normally does not permit activation of *comX* transcription by exogenously supplied XIP. The finding that this activation is unaffected by rapid replacement of the medium implies that the *comX* response in the *comS* overexpressing strain was not due to accumulation of extracellular XIP (or ComS). Curiously however, this *comX* response did require an intact chromosomal *comS*: Figs. 2A and 2B show that very few Δ*comS* cells activated *comX*, even though they harbored the *comS* overexpression plasmid. Together with Fig. 1, these data show that the chromosomal *comS* plays a role in *comX* activation that is not fully complemented either by saturating concentrations of exogenous XIP or by endogenous overproduction of ComS.

**Fig. 2:**
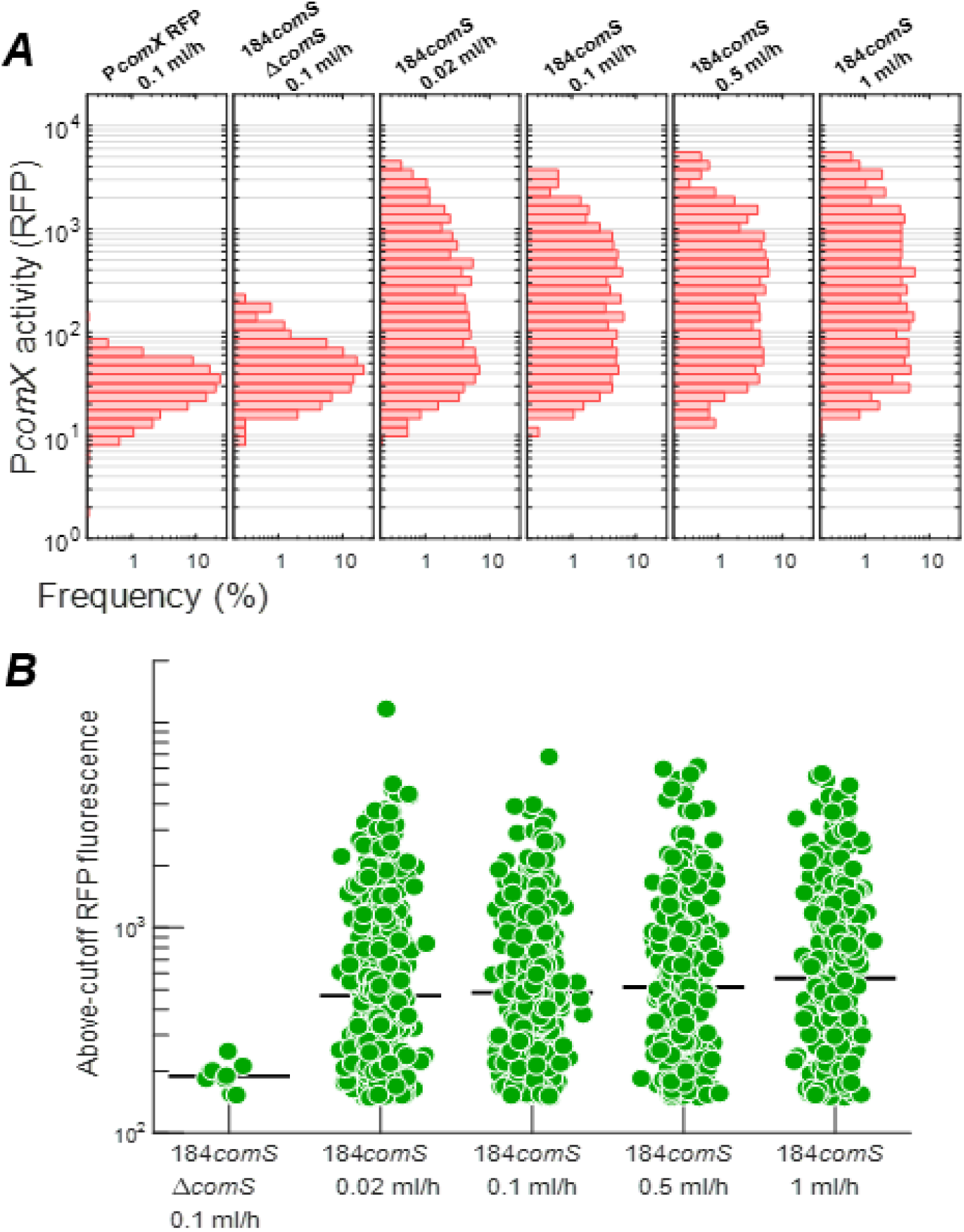
Activation of ComX in a ComS-overexpressing strain is independent of rate of medium replacement. P*comX-rfp* reporter activity is shown in cells growing in microfluidic chambers supplied with continuously flowing fresh complex medium (BHI). (A) Histograms of individual cell P*comX*-*rfp* reporter fluorescence: (Leftmost column) wild-type background (negative control) with flow at 0.1 ml h^-1^; (Second column) ComS-overexpressing 184*comS* Δ*comS* background at 0.1 ml h^-1^; (Columns 3-5) ComS-overexpressing (184*comS*) background at 0.02 ml h^-1^, 0.1 ml h^-1^, 0.5 ml h^-1^ and 1 ml h^-1^. (At a flow rate of 1 ml h^-1^ the medium in each flow chamber is replaced every 6 s.) (B) RFP fluorescence of cells that exceeded the wild type (negative control) red fluorescence in column 1 of (A). The black bar indicates the median of data in each channel: (Leftmost column) 184*comS* Δ*comS* background; (Columns 2-5) ComS-overexpressing (184*comS*) background. All RFP measurements were made 4 hours after addition of chloramphenicol to the cultures.

### The presence of *comS* in the chromosome affects *comS* mRNA levels

We used RT-qPCR to measure *comS, comR* and *comX* transcript copy numbers in mid-exponential phase cultures of the ComS overexpression, *comS* deletion, and UA159 strains (Fig. S1). Transcription of *comS* (Fig. S1A) was significantly increased in cells treated with exogenously added XIP in defined medium, compared to controls lacking XIP or growing in complex medium. In addition, cells overexpressing *comS* in a wild-type genetic background (184*comS*) and growing in complex medium had *comS* transcript levels that were significantly higher than a *comS* overexpressing strain that lacked the chromosomal copy of *comS* (184*comS* Δ*comS*). Levels of *comR* transcripts showed no change across strains or conditions (Fig. S1C).

### Population density of *comS* overexpressing cells does not determine the *comX* response

To further test whether *comS-*overexpressing cells release extracellular XIP, we measured the effect of *comS*-overexpressing (sender) populations on *comX* activation in *comS*-deficient (receiver) cells growing in coculture. We mixed sender (184*comS* P*comX-rfp*) and receiver (P*comX-gfp* Δ*comS*) cultures in different ratios and loaded them into microfluidic chambers containing static medium without exogenous XIP. We anticipated that, if senders released XIP or ComS into the extracellular medium, both senders (RFP reporter) and receivers (GFP reporter) would respond by activating *comX*, and that the average activation would increase with the ratio of senders to receivers. We used defined medium (FMC) to facilitate import of ComS/XIP if present.

We analyzed the green and red fluorescence of the co-cultures to generate histograms of individual cell fluorescence that reveal both the receiver (green) and sender (red) *comX* response, shown in Figs. 3A and 3B respectively. Representative microscopy images are shown in Figs. 3C-3H. In a control chamber containing only receiver cells exposed to 50 nM XIP, the GFP fluorescence was enhanced, but RFP fluorescence remained at the baseline level (Fig. 3D and second column in Figs. 3A, 3B). Similarly, a control chamber containing only sender cells showed enhanced RFP fluorescence, but only baseline GFP fluorescence (Fig. 3H and rightmost column in Figs. 3A, 3B). As expected, the median RFP fluorescence of the co-cultures (Fig. 3B, 3F-3H) increased at high sender/receiver ratios. The median RFP fluorescence of cells in the co-culture increased approximately in proportion to the density of senders, as expected if each sender activated its own *comX*. RFP fluorescence was constant over a period of four hours (Fig. S2).

**Fig. 3:**
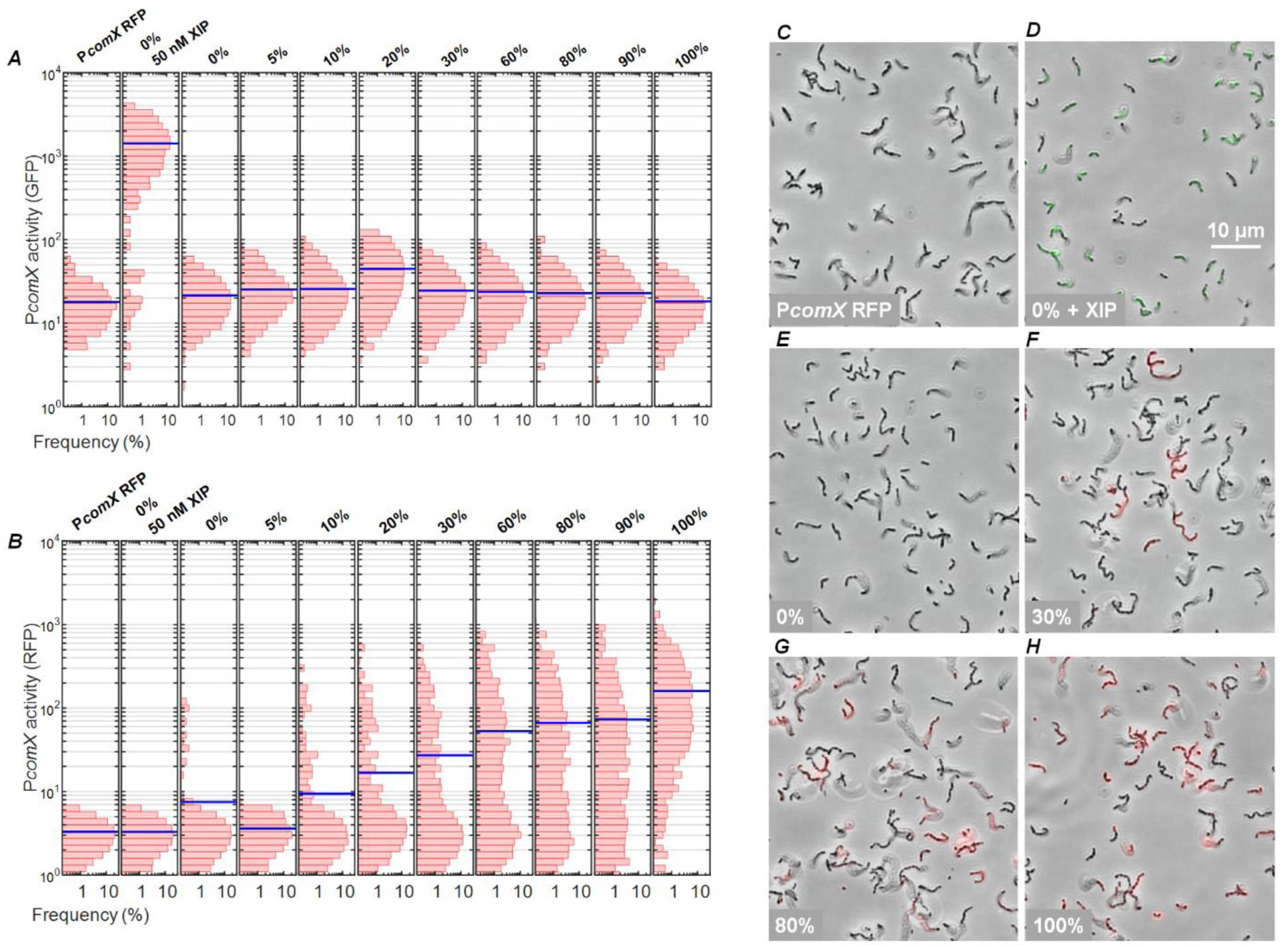
ComS overexpresser does not induce *comX* of Δ*comS* strain in coculture. Histograms of (A) GFP and (B) RFP fluorescence of individual cells in cocultures of sender (184*comS* P*comX-rfp*) and receiver (P*comX-gfp* Δ*comS*) strains. Strains were grown to equal optical density, mixed in varying proportion, and then incubated in microfluidic chambers containing stationary defined medium. Blue lines indicate population medians. Panels (A) and (B) show fluorescence of: (Leftmost column) UA159 background strain containing P*comX rfp* reporter, without added XIP (negative control); (Second column) P*comX-gfp* Δ*comS* (receiver) with added 50 nM XIP (positive control); (Columns 3-11) Cocultures of sender and receiver, with columns labeled by percentage by volume of 184*comS* (sender) culture in the initial preparation of the co-culture. (C-H) Phase contrast images of cocultures, overlaid with red and green fluorescence images: (C) P*comX-rfp* reporter in UA159 background, with no added XIP (negative control); (D) P*comX-gfp* Δ*comS* cells with 50 nM added XIP (positive control); (E) P*comX*-*gfp* Δ*comS* (receiver) alone, with 0% sender; (F)-(H) cocultures containing 30%, 80%, and 100% sender respectively.

However, when exogenous XIP was not provided, the GFP fluorescence of co-cultures showed no dependence on the sender/receiver ratio. Over the 0-100% range of co-culture ratios shown in Fig. 3A, the GFP fluorescence histograms did not exceed the baseline level of the negative control (Δ*comS* strain alone) in the third column of Fig. 3A. This GFP response did not change appreciably over a period of four hours (Fig. S2). Senders did not activate *comX* in the Δ*comS* receivers at any mixing ratio.

These data show that although overexpression of *comS* stimulates *comX* within individual cells, over the 4 h time scale of this experiment this activation did not cause extracellular, diffusible XIP to accumulate to levels sufficient to activate nearby Δ*comS* cells.

### Growth phase-dependent release of XIP

We previously showed that intercellular signaling by *S. mutans* ComRS can occur late in growth, but is impeded by deletion of the *atlA* gene, which encodes a major autolysin (26, 30). Loss of AtlA inhibits cell lysis, which appears to occur primarily in stationary phase. We therefore tested whether competence signaling from sender (*comS* overexpressing) to receiver (Δ*comS*) cells can be observed in the later phases of growth. We prepared sender/receiver co-cultures in different ratios in defined medium (to favor import of extracellular XIP if present). Because low pH suppresses the *comX* response, we adjusted the pH during the experiment to ensure that cells would remain responsive to *comX*-activating signals if present (9, 13). Every 2 h, the pH of the cultures was adjusted to 7.0 by addition of 2 N NaOH, the OD_600_ was recorded, and an aliquot of the culture was collected for fluorescence imaging of *comX* promoter activity. The GFP fluorescence histograms of Fig. 4A show that *comX* expression in the Δ*comS* strain increased very slightly at 12 h in comparison to expression at 2 h or 8 h. This increase was more pronounced at higher ratios of sender to receiver cells, consistent with some release of XIP from lysing senders late in the growth phase. The histograms of Fig. 4B, like those of Fig. 3B, show a strong RFP response (consistent with autoactivation of senders) with a slightly stronger response at earlier times.

**Fig. 4:**
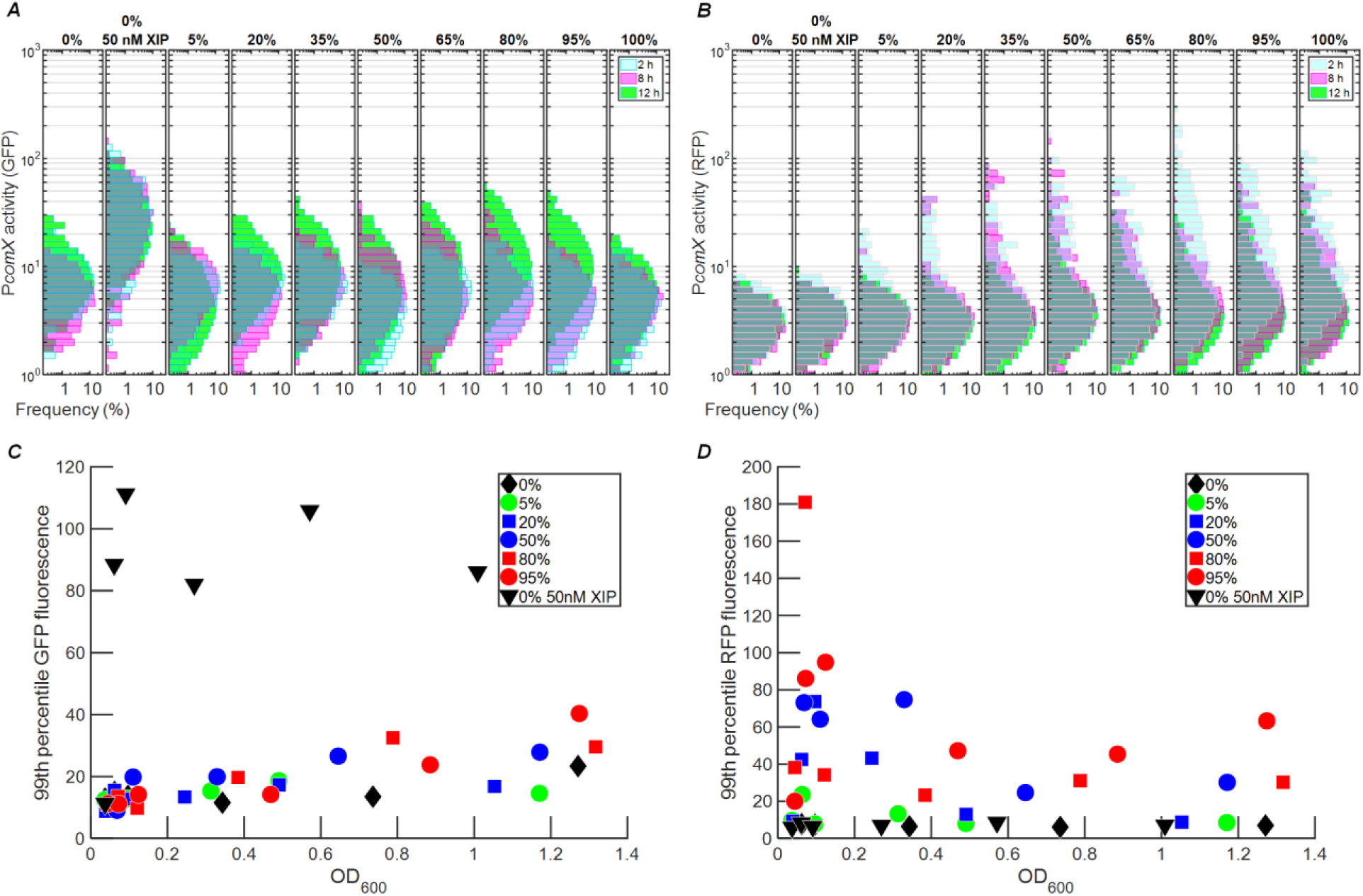
Evidence for release of XIP in cocultures late in growth. Histograms of (A) GFP and (B) RFP fluorescence of individual cells in cocultures of receiver (P*comX*-*gfp* Δ*comS*) and sender (P*comX*-*rfp* 184*comS*) strains, following different incubation periods. Labels at top indicate the volume fraction comprised by the sender strain in the preparation of the coculture. Histogram colors indicate incubation times: 2 h (cyan), 8 h (magenta), 12 h (green). The lower panels show the 99th percentile of the individual cell GFP (C) and RFP (D) fluorescence observed in the culture, versus the cultureOD_600_. Exogenous XIP was not added, except in the positive (sender-only) control sample indicated by the inverted triangles in (C) and (D).

The median GFP and RFP signals in the above histograms do not shift dramatically with either time or co-culture ratio. However, the histograms in Fig. 4A suggest moderate, density-dependent increases in receiver (green) fluorescence at 12 h. Figs. 4C and 4D highlight these changes by showing the value of the 99^th^ percentile of red and green fluorescence respectively in the cultures, versus optical density. In Fig. 4C, the GFP fluorescence of the most active receivers increases slightly at higher OD_600_ values, a change that is slightly more pronounced at higher sender:receiver ratios. By contrast, Fig. 4D shows no strong trend in the RFP fluorescence of the 99^th^ percentile of senders, versus OD_600_. None of the co-cultures exhibited as strong a GFP response as the positive control (receiver + 50 nM synthetic XIP, Fig. 4C), indicating that even 12 h growth did not lead to an extracellular XIP accumulation as large as 50 nM. Overall these data are consistent with robust self-activation of the *comS* overexpressing strain, accompanied by a modest release of XIP (or ComS) to the extracellular medium during the late stages of the growth curve.

### Binding *in vitro* of ComR to the *comS* and *comX* promoters

The observation of *comX* activation in the overexpressing (sender) strain, without significant accumulation of XIP in the medium, implies that activation of *comX* does not require export and reimport of XIP or ComS, if the chromosomal copy of *comS* is intact. To test whether endogenously produced ComS, acting intracellularly, is sufficient to activate *comX*, we tested whether unprocessed ComS could enable the binding of ComR to the *comS* and *comX* promoter regions *in vitro*. Fig. 5A shows a fluorescence polarization assay with purified recombinant ComR, synthetic XIP or ComS, and a fluorescently labeled DNA oligomer corresponding to the *S. mutans comX* promoter region containing the ComR binding site. The assay was performed in the presence and absence of excess (10 μM) ComS or XIP. Because of the excess of peptide (10 μM) relative to fluorescent DNA probe (1 nM), the probe polarization depends primarily on the concentration of ComR added. Fig. 5A shows that in the absence of XIP or ComS, ComR caused a weak rise in the fluorescence polarization of the DNA oligomer, indicating relatively weak affinity, as observed for ComR of other streptococci (20). However, in the presence of ComS or XIP the binding isotherm saturated at lower ComR concentrations, indicating formation of a complex with higher affinity for the *comX* promoter. Histidine tagging of ComR was found to reduce this affinity, as shown in Fig. S3.

**Fig. 5:**
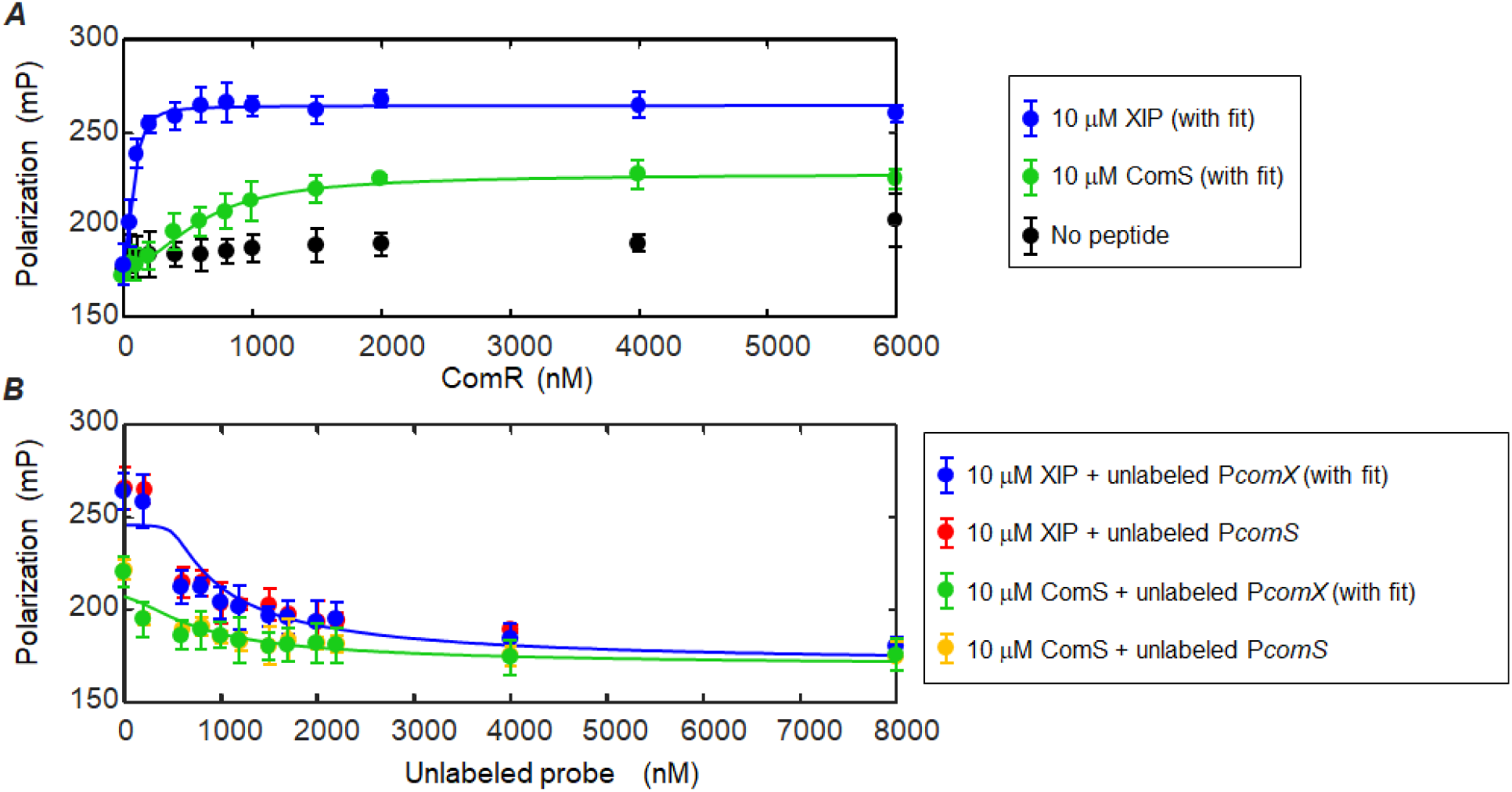
ComS and XIP interact with ComR to bind the *comX* promoter. Fluorescence polarization assay testing interaction of ComS and XIP peptides with ComR and the P*comX* transcriptional activation site. The DNA probe is labeled with a Bodipy FL-X fluorophore. (A) Titration of ComR into a solution containing 10 μM of XIP (blue) or full length ComS (green), labeled DNA probe (1 nM), and 0.05 mg ml^-1^ salmon DNA. Negative control (black) contains no ComS or XIP peptide. (B) Competition assay in which unlabeled promoter sequence DNA was titrated into a solution containing ComR (1.5 μM), fluorescent DNA probe (1 nM) and peptide (either ComS or XIP, 10 μM). Unlabeled P*comX* DNA was used with XIP (blue) and ComS (green). An unlabeled P*comS* probe was also tested for its ability to compete with the fluorescent P*comX* probe in the presence of XIP (red) and ComS (gold). Solid curves indicate binding and competition behavior predicted by the two step model described in *Methods*, in which peptide (ComS or XIP) first forms a multimeric complex with ComR (*k*_*1*_, *n*), and a single copy of this complex binds to the (labeled or unlabeled) P*comX* DNA (*k*_*2*_). For ComS binding/competition (green), the curves represent *k*_*1*_ = 3.2 μM, *k*_*2*_ = 2.2 nM, *n* = 2.5. For XIP binding/competition (blue), the curves represent *k*_*1*_ = 7.3 μM, *k*_*2*_ = 33 nM, *n* = 2.4.

Fig. 5B shows a competition assay in which unlabeled (‘cold’) P*comX* and P*comS* DNA oligomers having the same stem-loop structure as the labeled probe were titrated into samples containing 1.5 μM ComR, 10 μM ComS or XIP, and the labeled DNA (1 nM). The systematic decrease in polarization is consistent with competition for ComR. The unlabeled P*comS* and P*comX* probes, which differ by 3 bases, appear to have identical affinity for ComR.

Fig. 5A confirms a ComR-dependent interaction between ComS and the DNA probe, although this interaction is weaker than that of XIP. It also shows that, at saturating concentrations of ComR, ComS elicits only half of the total fluorescence polarization that results from an equivalent concentration of XIP. This difference may suggest that ComS and XIP induce a qualitatively different interaction between the ComR/peptide complexes and the DNA probe. The solid curves in Fig. 5A and 5B are calculated from a two-step, cooperative binding model (*Methods*) in which the dissociation constants for the ComR/peptide complex (*k*_*1*_) and the complex/promoter (*k*_*2*_), as well as the order of multimerization *n* of the ComR/peptide complex, are variables. Although the data clearly indicate that both ComS and XIP interact with ComR to bind the DNA probe, they do not permit a precise determination of the ComS and XIP interaction parameters. As discussed in *Methods*, the data are consistent with a range of parameter values, corresponding roughly to micromolar *k*_*1*_ and nanomolar *k*_*2*_ for both ComS and XIP, and similar cooperativity *n* for both peptides. The curves in Fig. 5 show the model with a roughly two fold difference in the ComR dissociation constants for ComS (*k*_*1*_ = 3.2 μM, *n* = 2.5) and XIP (*k*_*1*_ = 7.3 μM, *n* = 2.4).

### ComRS can drive expression from the *comX* promoter in *E. coli*

As discussed above, the OppA permease, which imports extracellular XIP, is not required for CSP-driven stimulation of ComRS and *comX*. In addition, the export of XIP is accomplished by the non-specific mechanism of lysis. To test whether the ComRS system requires dedicated mechanisms for processing of ComS to XIP, we constructed a dual plasmid system that placed IPTG-inducible *comS* and *comR* into *E. coli* cells carrying a P*comX-gfp* reporter. Fig. 6 shows the GFP activity of *E. coli* BL21(DE3) cells possessing the pDL278 P*comX*-*gfp* fusion and a pACYC-Duet1 that was either (*a*) empty plasmid or else carried inducible (*b*) *comS*, (*c*) *comR*, or (*d*) *comR* + *comS* (Fig. 6A). Fig. 6B shows GFP activity of the control strains (a-c) in response to added IPTG. Following induction by IPTG, the *comS* and empty vector controls show only baseline GFP expression (Fig. 6B). However, *comR* alone was able to elicit some expression from the *comX* reporter, consistent with the weak DNA binding seen at high ComR concentrations in Fig. 5. When both *comR* and *comS* were present (Fig. 6C), the median *comX* expression was roughly 2-fold greater than with ComR alone at 1 mM IPTG, and roughly 3-fold higher at 300 μM IPTG. These data show that endogenously produced, unprocessed ComS is sufficient to enhance ComR-induced expression from the *comX* promoter.

**Fig. 6:**
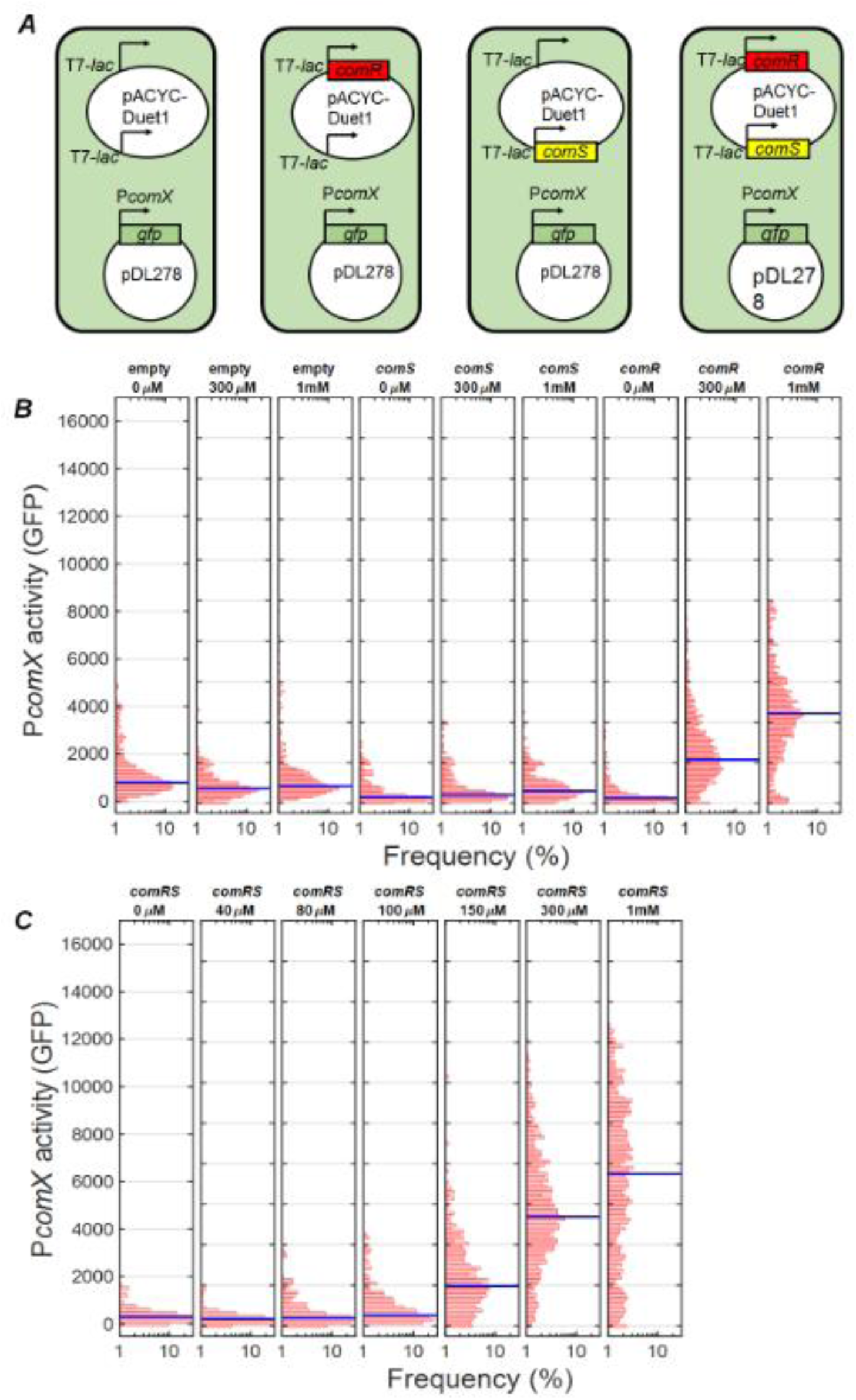
Activation of P*comX* by inducible *comRS* in *E. coli*. Activity of a P*comX-gfp* reporter within individual *E.coli* cells measured by fluorescence microscopy. Strains (A) harbored a two-plasmid system that included the P*comX-gfp* reporter as well as different combinations of *comR* and *comS* under control of the T7-*lac* promoter. (B) GFP fluorescence of controls containing the pACYC-Duet1 empty vector, *comR* alone or *comS* alone after 3 h of growth with shaking, following addition of 0, 300 μM and 1 mM IPTG. (C) GFP fluorescence of the strain possessing both *comR* and *comS* after 3 h of growth with shaking in varying IPTG concentrations. The blue bar represents the median cell fluorescence.

### Data are consistent with an intracellular feedback loop in *comS* transcription

The reduced *comX* response in *comS* deletion strains in Fig. 1A and Fig. 2A shows that an intact *comS* has a stimulatory effect on *comX* expression that is not entirely matched either by exogenous XIP or by overexpression of ComS from a plasmid. One explanation for this effect is transcriptional positive feedback, in which endogenously produced ComS retained within the cell interacts with ComR to stimulate transcription from *comS* and *comX* (Fig. 7A). Import of extracellular XIP would be expected to stimulate such feedback in cells carrying an intact *comS*, potentially accounting for the stronger *comX* response to XIP that is observed in the wild type versus Δ*comS* background.

**Fig. 7:**
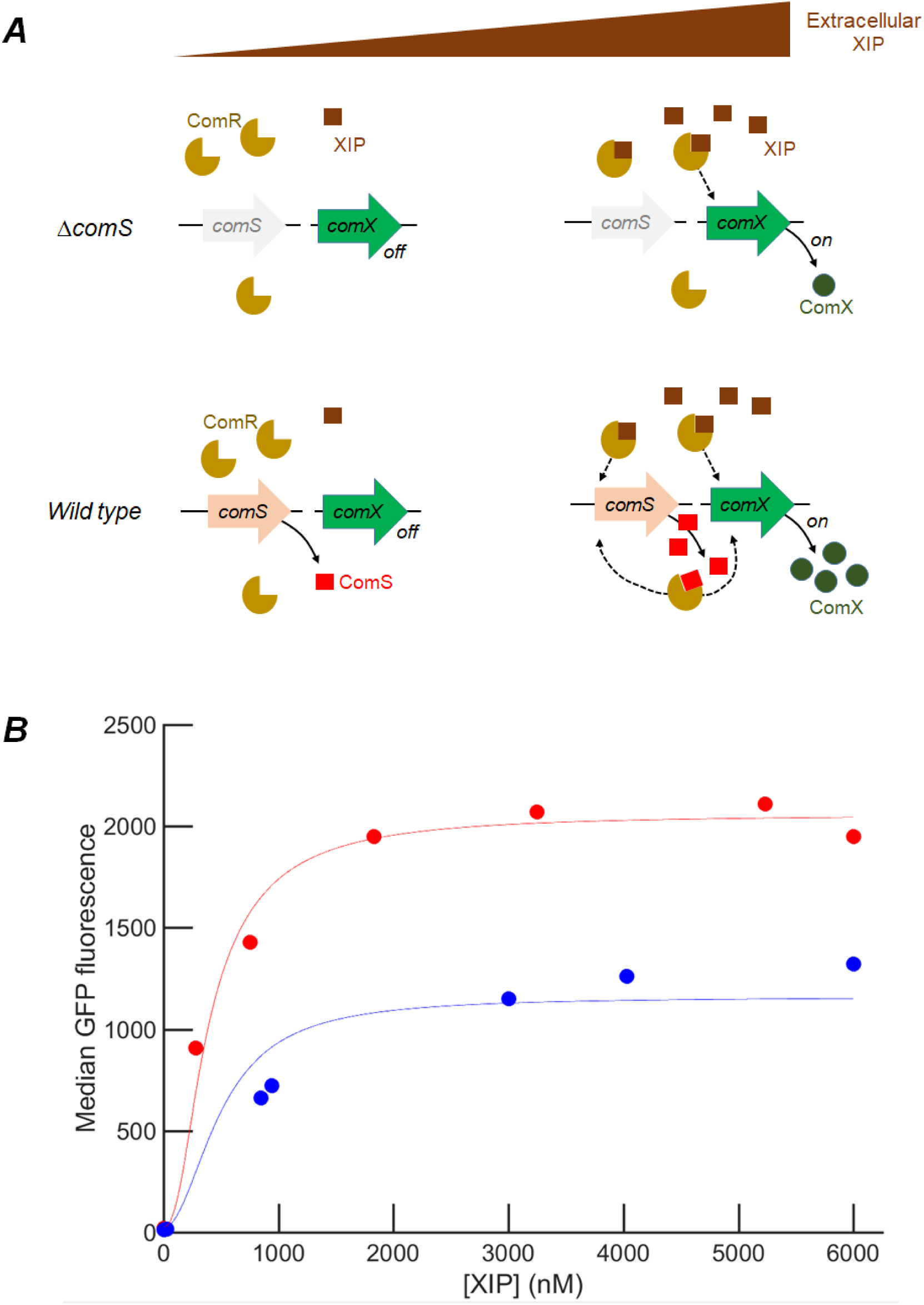
proposed model for *comRS* feedback regulation of *comX*. Model for *comS*-feedback enhanced activation of P*comX-gfp* reporter by exogenous synthetic XIP. (A) Illustration of the feedback model and the role of *comS* and extracellular XIP. ComS and XIP both interact with ComR to activate transcription of *comS* and *comX.* At low concentrations of extracellular XIP, *comS* expression is very low and *comX* is not expressed. At high concentrations of extracellular XIP, XIP is imported efficiently by Opp and interacts with ComR to drive expression of both *comS* and *comR*. Endogenously produced ComS is not readily exported in the absence of lysis, and so intracellular accumulation of ComS drives elevated *comS* and *comX* expression. Consequently *comX* expression at any given XIP concentration is boosted by *comS* feedback. Cells lacking native *comS* can respond to synthetic XIP but cannot activate *comX* to the same level as wild type. The figure describes behavior in defined medium; in complex medium extracellular XIP is not imported (11). (B) Comparison of model simulation with data. Red circles indicate median P*comX-gfp* fluorescence of the UA159 background strain supplied with synthetic XIP in microfluidic flow; blue circles indicate the median P*comX-gfp* fluorescence of the Δ*comS* background. Solid curves represent calculated values from a fit in which 11 parameters were fit to the microfluidic data, as described in *Methods.* The model relates the predicted ComX concentration to the median GFP fluorescence by an offset and scale factor.

We constructed a mathematical model of this mechanism as described in the *Methods* and the Supplemental Information. The model addresses conditions in a defined medium where extracellular XIP can be imported to the cell, and early growth where lysis-driven release of ComS or XIP is not significant. Under these conditions the model assumes that (I) extracellular XIP is imported to the cell but (II) neither ComS nor XIP is exported. Further (III) both the ComS and XIP complexes of ComR are able to elicit transcription of *comS* and *comX*, although not necessarily with equal transcription rates or degree of multimerization. Assumption (III) is motivated by data indicating that ComR binding does not absolutely require processing of ComS to XIP, by differences in the saturating fluorescence polarization achieved with ComS and XIP in Fig. 5A, and by the different saturating *comX* response (to XIP) of the Δ*comS* and UA159 backgrounds in Fig. 1A. In parametrizing the model we allow a different order of multimerization in the ComR/peptide complex that binds DNA: *n* = 1 (ComS) or *n* = 2 (XIP) (see *Discussion*). In addition the maximal rate of transcription (ComX production) is permitted to depend on whether a ComR/XIP or a ComR/ComS complex is bound to the promoter.

Fig. 7B shows the results of solving the steady state dynamical equations of the model with parameter values that best reproduce the data (*Methods*). If only XIP (but not full-length ComS) interacts with ComR to activate *comX*, the model gives identical *comX* activation in both the Δ*comS* and wild-type backgrounds, in contradiction of Fig. 1A. However if both the ComR/XIP and ComR/ComS complexes are permitted to induce transcription, although with different maximal rates, the model reproduces the different *comX* expression in the Δ*comS* and wild-type backgrounds. Parameter values (Table S2) were found to preserve the relative orders of magnitude of the dissociation constants of the transcriptional activators found in Fig. 5, with the best fit giving a transcription rate that is between 4-fold and 200-fold greater for the ComS-ComR bound promoter than for XIP-ComR activation of the gene.

## Discussion

The ComRS system found in mutans, salivarius, pyogenic and bovis streptococci has been described alternatively as a quorum sensing (19) or timing (25) mechanism that directly controls *comX*, the master regulator of genetic competence. The ComS-derived peptide XIP is readily imported by *S. mutans* in defined growth medium, where it induces transformability with high efficiency. Key evidence supporting an intercellular signaling role for XIP includes the detection by LC-MS/MS spectroscopy of XIP in supernates of *S. mutans* that were grown to high cell densities (22, 23). In addition, filtrates of *S. mutans* cultures grown to high density induced P*comX* activity in reporter strains (21), indicating the presence of an active competence signal in the extracellular medium. A recent co-culture study verified that XIP is freely diffusible in aqueous media and that ComS-overexpressing senders are able to activate *comX* in nearby Δ*comS* receiver mutants, with no cell-cell contact being required (26). However, deletion of *atlA*, which encodes a surface-localized protein associated with envelope biogenesis and autolysis (30), suppressed this intercellular signaling (26). Taken together these data indicate that the *S. mutans* ComRS system can provide intercellular competence signaling when autolysis releases sufficient concentrations of ComS or XIP.

We have previously argued that the bimodal expression of *comX* under stimulation by CSP is a signature that *comX* is controlled by an intracellular transcriptional feedback system: endogenously produced ComS (or its product XIP) accumulates within some cells and then interacts with ComR to activate *comX.* Such feedback mechanisms are often a cause of bimodality in bacterial gene expression, including in other competence pathways (31, 32). The lack of an established mechanism for processing and export of ComS, the sensitivity of *comX* bimodality to endogenous production (via chromosomal *comS)* of XIP, and the fact that XIP import via Opp is not required for the bimodal behavior, together suggest that purely intracellular signaling in *comRS* may be possible, at least under some environmental conditions. Such a mechanism would potentially allow individual *S. mutans* within a population to differ in competence behavior, as observed experimentally (14).

The present data support this model by providing several lines of evidence that the chromosomal *comS* plays a role in ComRS activation of *comX*, regardless of other XIP or ComS that may be supplied. First, although the response of *comX* is different in complex medium supplemented with CSP than in defined medium supplemented with XIP, in both cases the behavior of *comX* is affected by the presence of the intact chromosomal *comS* under the control of its cognate promoter. In complex medium, *comS* is required in order for CSP to elicit any *comX* response; in defined media the deletion of *comS* reduces the *comX* response (both average and variance) to exogenous XIP, and raises the threshold XIP concentration for a response. Further, even if it harbors a *comS* overexpression plasmid, a *comS* deletion strain expresses *comX* much more weakly than does a *comS-*overexpressing strain that retains its chromosomal *comS*. These data show that the cell’s own native regulation of *comS* affects its activation of *comX*, independent of whether it overproduces ComS from a plasmid or imports exogenous XIP via Opp.

Our data also show that even in complex growth medium, which is known to inhibit the uptake of extracellular XIP, ComS-overexpressing (sender) cells activate their own *comX.* This autoactivation is unaffected by very rapid exchange (by flow) of the medium, strongly suggesting that *comX* activation in these cells does not require accumulation of extracellular XIP. However, this autoactivation requires a native *comS* in addition to the overexpression plasmid. In addition, even in the ComS overexpression strain, *comS* mRNA levels are significantly higher when the native *comS* is present, but *comR* mRNA levels are unaffected (Fig. S1). These behaviors are consistent with positive feedback where imported XIP or plasmid-generated ComS enable internal amplification of *comS* expression.

Finally, the data show that *comS*-overexpressing cells fail to stimulate Δ*comS* cells (receivers) in co-cultures in defined medium conditions, which are favorable for import of XIP. As the Δ*comS* receivers do respond to exogenously added XIP, these data indicate that the overexpressing cells activate their own *comX* without releasing significant XIP to the medium. The weak intercellular signaling that is observed in co-cultures grown to late growth phases is consistent with eventual lysis of sender cells, possibly linked to autoactivation of the lytic pathway driven by ComX and ComDE.

The finding that diffusive signaling by ComS or XIP between *S. mutans* cells is inefficient or lacks spatial range is consistent with the conclusion reached by Gardan *et al*. using *S. thermophilus* (25). Those authors found that the type I ComS peptide of *S. thermophilus* was not secreted at detectable levels in a strain that produced it naturally, although an overproducing strain did generate detectable ComS in the medium. They argued that ComS does not diffuse through or accumulate in the medium, although it may be able to signal between cells that are in physical contact. This proximity model for ComS resembles a “self-sensing” quorum system (33) in which the secreted signal is retained at elevated concentrations in the immediate surroundings of the cell, possibly associated with the cell surface, so that the cell responds somewhat more strongly to its own secreted signal than to that of the rest of the population.

Our observations suggest more strongly that intercellular quorum signaling through ComS or XIP is not essential to ComRS control of *comX* in *S. mutans*, under the conditions examined. A more essential component is the dynamics of the cell’s own *comS* transcription. A plausible mechanism for the bimodal response of *S. mutans comX* to CSP stimulation is therefore that CSP stimulates the bistable *comRS* feedback loop by facilitating, through an indirect mechanism, the positive feedback. It would be sufficient for example to inhibit degradation of endogenous ComS, which depending on basal transcription levels would trigger *comX* activation in at least some cells, leading to the bimodal distribution of *comX* activity (11). Notably our data show that overexpression of *comS* also leads to heterogeneous *comX* activity, suggesting that it plays a role similar to exogenous CSP by facilitating *comS* autofeedback.

As it is unknown in *S. mutans* whether ComS is processed to XIP inside the cell, either XIP or ComS could potentially act as the intracellular feedback signal. Although *S. mutans* competence was shown to be unresponsive to exogenous full-length ComS (19), this finding may reflect either selectivity by ComR or simply inefficient import of full length ComS by Opp. ComS is significantly larger (17 residues) than peptides that are typically transported by ABC transporters. Shanker *et. al.* (34) found that *S. mutans* ComR is unresponsive to the ComS peptides produced by other streptococcal species, although an eight residue XIP (ComS_10-17_) did interact effectively with ComR to bind the *comS* and *comX* promoters (20). Our fluorescence polarization data confirm that both ComS and XIP can interact with ComR to bind the *comX* and *comS* promoter regions. They also suggest that ComS and XIP may form ComR complexes of different degrees of multimerization, a difference that could have interesting consequences for the nonlinear dynamics of feedback regulation. Our mathematical model for transcriptional autofeedback in the *comRS* system incorporates the data by assuming that endogenously produced ComS is not released to the environment, although extracellular XIP is imported and supplements the endogenous ComS in interacting with ComR. Unless *E. coli* is equipped to process ComS to XIP, or perhaps that ComS cleaves autocatalytically, our finding that *E. coli* carrying inducible *comR* and *comS* can activate a *comX* reporter plasmid is further evidence that processing of ComS to XIP is not absolutely required.

Other studies show some precedent for such regulation. Structural studies in *S. pyogenes* have shown that some intracellular Rgg receptor proteins can bind pheromones that differ in length and sequence (35). Crystallographic structures of homologous ComR proteins (34, 36) show the SHP binding pocket of the ComR C-terminus to fall in the tetratricopeptide repeat domain that is responsible for multimerization, while the N-terminus helix-turn-helix structure binds DNA after an induced structural rearrangement. The location of the SHP binding pocket could allow the longer ComS to hinder multimerization when bound, resulting in a monomer binding to its target, while XIP does not. Fig. S3 shows preliminary evidence that the ComS N-terminus affects ComR binding in *S. mutans*. Neither ComS nor XIP could induce DNA binding by an N-terminally 6x histidine tagged ComR, whereas XIP (but not ComS) caused DNA binding activity in a C-terminally tagged ComR. These data indicate that steric effects around the SHP binding pocket may influence DNA binding affinities.

Positive feedback occurs in many quorum sensing systems as the accumulation of the chemical signal in the extracellular environment stimulates the cell to produce additional signal or its cognate receptor. For example, in *Vibrio fischeri* the C8 homoserine lactone autoinducer stimulates expression of *ainS*, which encodes the autoinducer synthase (37). In *Vibrio cholerae*, the CAI-1 signal stimulates production of its receptor CqsS (38). In these cases, the extracellular signal concentration, which is the positive feedback signal, is sensed by large numbers of cells, and so the population responds homogenously. However, if an individual cell responds preferentially to its own signal production, then the feedback signal is specific to the individual cell and the behavior is qualitatively different. Individual feedback can convert a graded (or unimodal) population response to a switched or bimodal response (39). Depending on parameters such as the rate of signal production, noise levels or the cell density, the response of the cells may then span a range from strongly social or quorum behavior to purely autocrine or self-sensing (33) behavior in which cells respond independently and the population becomes heterogeneous (40). Synthetic biology has exploited this phenomenon in several bacterial quorum sensing systems to amplify the cell’s sensitivity to an exogenous signal. This can lower the quorum circuit’s threshold sensitivity to the signal, and it can also enhance the amplitude of the cell’s full response to that signal. Figs. 1A and 1B suggest that the chromosomal *comS* in *S. mutans* roughly doubles the amplitude of *comX* response and lowers the XIP sensitivity threshold roughly two-fold. This amplification is comparable to what was accomplished in engineered, synthetic systems (41, 42).

As a result the ComRS system may have two modes of function in *S. mutans*. At low population densities, during early growth, ComRS operates through intracellular feedback, leading to population bimodality in *comX* expression. Here, only a small subpopulation of cells activate the late competence genes. However, in later growth phases or in mature biofilms, stress mechanisms that drive autolysis allow the release of XIP, providing a diffusible signal that is detected by other cells and amplified through the internal feedback mechanism to elicit a strong competence response. In this sense, XIP may serve to broadcast localized stress conditions, stimulating *S. mutans* to scavenge DNA resources opportunistically from nearby lysing cells (43, 44).

## Materials and methods

### Strains and growth conditions

*S. mutans* wild-type strain UA159 and mutant reporting/gene deletion strains from glycerol freezer stock were grown in BBL BHI (Becton, Dickinson and co.) at 37°C in 5% CO_2_ overnight. Antibiotics were used at the following concentrations where resistance is indicated in Table 1: erythromycin (10 μg ml^-1^), kanamycin (1 mg ml^-1^), spectinomycin (1 mg ml^-1^). For experiments in defined medium, strains were washed twice by centrifugation, removal of supernatant and re-suspension in the defined medium FMC (45). These were then diluted 20-fold into fresh FMC and allowed to grow in the same incubator conditions until at optical density at 600 nm (OD_600_) of 0.1. Synthetic XIP (sequence GLDWWSL) was synthesized and purified to 98% purity by NeoBioSci (Cambridge, MA).

**Table 1:** list of strains and plasmids used.

| Strain or plasmid | Characteristics* | Source or reference |
| --- | --- | --- |
| ***S. mutans* strains** |  |  |
| *PcomX-gfp* (plasmid) | UA159 harboring *PcomX* GFP promoter fusion on plasmid pDL278. | (11) |
| *PcomX-rfp* | UA159 harboring *PcomX* dsRed RFP promoter fusion on plasmid pDL278. | (26) |
| *PcomX-gfp ΔcomS* | UA159 *comS* gene replaced with a non-polar erythromycin resistance cassette. Harboring *PcomX* GFP promoter fusion on plasmid pDL278. | (11) |
| 184*comS PcomX-rfp* | UA159 harboring pIB184*comS* and *PcomX* dsRed RFP promoter fusion. | (26) |
| 184*comS PcomX-rfp ΔcomS* | UA159 harboring pIB184*comS* and *PcomX* dsRed RFP promoter fusion. *comS* disrupted by | This study. |
|  | point mutation in start codon (ATG to AAG). |  |
| ***E. coli* strains** |  |  |
| BL21(DE3) | Used for recombinant protein expression | New England Biolabs, MA |
| 10-beta | Used for propagating plasmids during cloning | New England Biolabs, MA |
| SAMU215 | BL21(DE3) carrying *PcomX-gfp* on pDL27 | This study |
| SAMU216 | SAMU215 carrying pACYC-Duet1 empty vector | This study |
| SAMU217 | SAMU215 carrying pACYC-Duet1 plasmid with *comR* on multiple cloning site (MCS) I | This study |
| SAMU218 | SAMU215 carrying pACYC-Duet1 plasmid with *comS* on MCS II | This study |
| SAMU219 | SAMU215 carrying pACYC-Duet1 plasmid with *comR* on MCS I and *comS* on MCS II | This study |

| Plasmids |  |  |
| --- | --- | --- |
| pIB184 | Shuttle expression plasmid with the P23 constitutive promoter, Em<sup>R</sup> | (29) |
| pDL278 | *E. coli* – *Streptococcus* shuttle vector, Sp<sup>R</sup> | (46) |
| pET45b(+)*his-comR*<sub>UA159</sub> | pET45b(+) derivative containing the translational fusion P<sub>T7lac</sub>-6*xhis-comR*<sub>UA159</sub>, Ap<sup>r</sup> | This study |
| pACYC-Duet 1 | T7-*lac* inducible expression vector for co-expression of two proteins, Cm<sup>R</sup>. | Millipore-Sigma, MA |

\*Em = erythromycin, Sp = spectinomycin, Ap = ampicillin, Cm = chloramphenicol.

*E. coli* strains were grown in LB at 37 °C shaking in an aerobic incubator overnight. Antibiotics were used at the following concentrations where resistance is indicated: ampicillin (10 μg ml^-1^), chloramphenicol (34 μg ml^-1^). For recombinant ComR expression, the next day the overnight cultures were diluted 100-fold into LB containing ampicillin at the indicated concentration and grown under the same incubator conditions as overnight. For recombinant *comRS* expression inducing P*comX*-*gfp*, cells were diluted 100-fold into LB containing chloramphenicol and spectinomycin and grown shaking at 37 °C to an OD_600_ of 0.1, at which point IPTG was added and the cells grown shaking at 30 °C until imaging.

### Construction of *comS* point mutant

The start codon of the *comS* gene was mutated from ATG to AAG. The mutation was introduced directly into the chromosome by site-directed mutagenesis using a PCR product generated by overlap extension PCR (47). Potential mutants were screened using mismatch amplification mutation analysis (MAMA) PCR (48), as previously described (49, 50). The point mutation was confirmed by PCR and sequencing to ensure that no further mutations were introduced into the *comS* gene and its flanking regions.

### Microfluidic experiments

Microfluidic experiments were performed using a seven-channel PDMS-cast mixing array device, with fluorescence imaging and single-cell image analysis as described previously (11, 51, 52). Cells carrying a P*comX-gfp* plasmid-based reporter were grown to OD_600_ 0.1 from dilution in FMC and sonicated briefly using a Fisher Scientific FB120 sonic dismembrator probe to split large chains. Sonicated cells were then loaded into the device through a syringe capped with a 5 μm filter to remove any remaining aggregations. FMC containing 1 mg ml^-1^ spectinomycin and a XIP gradient produced from three inlets containing different concentrations of XIP (0 nM, 600 nM and 6 μM XIP inlets) passed through a mixing matrix was pumped through the cell chambers at a steady rate of 0.08 ml h^-1^ to create a constant, different XIP concentration in each cell chamber.

Gamma distributions (a two-parameter probability distribution describing the amount of protein produced in sequential transcription and translation steps) were fit to the single cell fluorescence distributions using Matlab to fit protein production to theoretical description (28). The fit was applied to cells fluorescing above an arbitrary cutoff of 40 units (around background) in order to prevent turned-off cells from skewing the distribution. Parameters were rounded to three significant figures and reported in Table S1.

### Flow rate dependence experiment

In order to measure the flow rate dependence of XIP signaling we loaded cells into a commercial six-channel microfluidic slide (IBIDI μ-slide VI, IBIDI GmbH). The six channels (*a-f*) contained respectively (*a*) a red fluorescent protein (dsRed) *comX* reporting strain (P*comX*-*rfp*) control channel flowing fresh BHI at 0.1 ml h^-1^; (*b*)-(*e*) four channels containing a *comS* overexpression strain 184*comS* P*comX-rfp* (*comS* on plasmid pIB184 under the strong constitutive P23 promoter) with BHI at different flow rates ranging from 0.02 ml h^-1^ up to 1ml h^-1^, and (*f*) a 184*comS* P*comX-rfp* strain with a point mutation disrupting the chromosomal *comS* gene (Δ*comS*) under flow at 0.1 ml h^-1^. After 2 hours the plain BHI supplied was replaced with BHI supplemented with 50 μg ml^-1^ chloramphenicol in order to halt further translation and allow any RFP in the cells to fold. This was supplied at 0.1 ml h^-1^ flow rate for all channels. Four hours (the maturation time of our RFP) after chloramphenicol addition final fluorescence images of the cultures were taken. Due to the bimodal *comX* activation in BHI, a fluorescence cutoff was set as the maximum RFP fluorescence observed in the PcomX-*rfp* negative control. Cells exhibiting RFP fluorescence above this level were collected in an array and the size of this sample as a percentage of the population as well as the median of the above-cutoff fluorescence reported.

### Channel co-culture experiment

We loaded co-cultures of a P*comX-gfp* Δ*comS* (responders) with the *comS* overexpressing strain 184*comS* P*comX*-*rfp* (senders) into two commercial microfluidic slides (IBIDI μ-slide VI) using static (not flowing) FMC medium and varying ratios of *comS* overproducers:Δ*comS* responders (percentage by volume of OD_600_ 0.1 cultures vortexed together). P*comX-rfp*, P*comX-gfp* Δ*comS*, P*comX-rfp* + 50 nM XIP and P*comX-gfp* Δ*comS* + 50 nM XIP were used as controls. The end ports of the channels were sealed with mineral oil to prevent drying of the medium in the channels. Images were taken as in the microfluidic experiments and analysis performed similarly. In the case of controls XIP was added to planktonic culture and the tube vortexed before pipetting into the slide. Because the population was heterogeneous in both fluorescent reporter type and *comX* expression, a fluorescence threshold was defined as the maximum RFP fluorescence observed in the P*comX*-*rfp* negative control as previously. The median of the RFP fluorescence observed above this cutoff in other samples was used as a measure of how strongly the red cells were activating *comX* as a function of their number density.

### OD dependence of co-culture response

For tests of growth-phase dependence of signaling, co-cultures similar to those in microfluidic channel slides were prepared. Overnight cultures were washed and diluted 40x into fresh FMC containing erythromycin (10 μg ml^-1^) and spectinomycin (1 mg ml^-1^). Once grown to OD_600_ 0.05, these were mixed in ratios varying from to 0% *comS* overexpressers to 100% overexpressers, defined by volume of *comS* overproducers added divided by the volume of the Δ*comS* culture added. Low initial cell densities were used to ensure that early, mid and late growth phases were probed for XIP release. Every two hours the OD_600_ of the culture and its pH were measured. The pH was corrected back to 7.0 using 2N sodium hydroxide if it had deviated below 6.5, in order to measure reaction to any XIP released at late times into the culture. RFP and GFP fluorescence were measured by pipetting a small amount of the culture onto a glass coverslip and analyzing single cells. 99^th^ percentile GFP fluorescence was then used to determine if XIP was being released to the *comS* mutants in an OD_600_-dependent manner.

### RT-qPCR measurement of *comS, comR* and *comX* transcripts

*S. mutans* cells were diluted 20-fold into BHI (P*comX-gfp* WT/BHI, 184*comS PcomX-rfp*, 184comS P*comX-rfp* ΔcomS samples) or FMC (P*comX-gfp* WT/FMC ± XIP, P*comX*-*gfp* Δ*comS* ± XIP samples, pIB184 / WT, pIB184ComS / WT). Where added, XIP was supplied at OD_600_ = 0.1. Cells were harvested at OD_600_ = 0.5 by centrifugation and re-suspended in RNA protectant buffer for 10 minutes. Samples were then centrifuged, the supernatant removed and the pellets frozen at −80 °C. RNA extraction was performed using the Qiagen RNEasy mini kit (Qiagen, USA). RNA sample concentration and purity were measured using a Thermo Scientific NanoDrop One Microvolume UV-Vis Spectrophotometer (Thermo Scientific, USA). 1 μg of RNA was then reverse transcribed to cDNA using the Bio-Rad iScript reverse transcription kit with random primers (Bio-Rad, USA). The qPCR was performed on a Bio-Rad CFX96 Real-Time System using Bio-Rad Sso Advanced Universal SYBR Green Supermix with a 50-fold dilution of the cDNA and 500 nM gene-specific primers. Sequences used for the primers are given in Table S3 (supplemental information). A standard curve across 8 orders of magnitude of transcript copies (from 10^8^ to 10^1^) was used to determine transcript count for each gene. For each sample the *comX, comR* and *comS* transcript counts were then normalized by the 16S rRNA count for the same sample. Fig. S1 shows the median of this ratio, with error bar lengths given by the range from second lowest to second highest ratio obtained.

### Fluorescence polarization

ComR protein was obtained by cloning the *comR* gene into the 6x-His tagged site on pET-45b(+) vector in *E. coli* 10-beta using standard PCR cloning methods. His-ComR was then expressed in *E. coli* BL21(DE3) by induction with 1mM IPTG at mid-exponential phase in LB. After 4 hours cells were lysed using lysozyme in B-PER lysis buffer (ThermoFisher). Protein was then purified from clarified lysate using Ni-NTA agarose affinity chromatography and the histidine tag cleaved using enterokinase max at 4°C (EKMax, Invitrogen). The resulting protein solution was dialyzed into PBS pH 7.4 for experimental use. Native ComR concentration was measured by the Pierce BCA assay (Thermo Scientific) and purity of the cleaved form verified by SDS-PAGE run against an uncleaved sample.

Fluorescence polarization assays were performed in a 96-well plate with black bottom and black sides in a Biotek Synergy 2 plate reader (Biotek Instruments inc.) in the polarization mode. A 5’ Bodipy FL-X labeled self-annealing stem-loop DNA strand with sequence corresponding to PcomX (sequence 5’-BODIPY FL-X - ATGGGACATTTATGTCCTGTCCCCCACAGGACATAAATGTCCCAT - 3’), synthesized by ThermoFisher) was used as the binding aptamer and a filter set with excitation 485 nm, emission 528 nm was used for fluorescence excitation. 1 nM labeled DNA probe was added to a reaction buffer previously described (53) supplemented with 1 mM EDTA and 0.05 mg ml-1 salmon DNA. ComR was titrated in concentration in this buffer alone, in the presence of 10 μM XIP or in the presence of 10 μM *comS*. The reactions were incubated at 37°C for 20 minutes before reading. Synthetic ComS (sequence MFSILTSILMGLDWWSL) for fluorescence polarization was synthesized and purified to 60% purity by Biomatik (Wilmington, DE).

Competing unlabeled probe assays were performed with 1.5 μM ComR in the same buffer containing 1 nM P*comX* fluorescent DNA and 10 μM SHP (either ComS or XIP). An unlabeled probe corresponding to either the P*comS* (sequence 5’ - ACG GGACATAAATGTCCTGTCCCCCACAGGACATTTATGTCCCGT - 3’), synthesized by Thermo Fisher) or the above P*comX* probe was titrated into this solution and the decreasing polarization plotted. Reactions were again incubated at 37°C for 20 minutes before polarization readings taken. In all FP experiments reading was performed three times on the same plate to estimate instrument error. The average polarization was used for plotting and analysis. Details and parameters of the two-step binding model for the FP data are given in the supporting information.

### ComRS expression in *E. coli*

A recombinant *comRS* system was produced in *E. coli* by inserting *comR* into multiple cloning site (MCS) I in the pACYC-Duet 1 vector and *comS* into MCS II by standard cloning methods. BL21(DE3) cells containing P*comX-gfp* on pDL278 were transformed with the resulting pACYC-Duet 1 *comRS* plasmid. Controls consisting of an empty pACYC-Duet 1 and the vector containing only *comR* at MCS I and only *comS* at MCS II were produced in the same way. Cells were diluted 100-fold from overnight into LB containing chloramphenicol and spectinomycin and grown shaking at 37 °C until they reached OD_600_ 0.1. At this point IPTG was added in varying amounts to induce expression of the T7-*lac* controlled recombinant system. Following addition of IPTG the cells were grown with shaking at 30 °C and GFP fluorescence was imaged after 3 h.

### Mathematical model of *comRS* control of *comX*

Deterministic modeling of *comX* activation by *comRS* was performed by least squares fitting a chemical equilibrium model to the microfluidic data from experiments for each of the wild type background P*comX* GFP strain and the Δ*comS* cells. Details of the model and the robustness analysis are in the supporting information, with parameter values given in Table S2.

## Acknowledgments

This work was supported by 1R01 DE023339 from the National Institute of Dental and Craniofacial Research. The authors thank Minjun Son, Chris Browngardt, Natalie Maricic, Lin Zeng, Hey Min Kim and Sang-Joon Ahn for helpful discussions, provision of mutant strains and advice on experimental procedures.

