## Supplementary Materials for "Intracellular signaling by the *comRS* system in *Streptococcus mutans* genetic competence" (from another version of this preprint: version 1, 2018-03-28)

List of supporting figures and tables:

- Figure S1: Response of cocultures is time-independent
- Figure S2: Effect of histidine tag on ComR binding of ComS and XIP
- Deterministic fit:  Equation system and calculated parameter values
- Table S1: Parameters for gamma distribution fits to single cell P*comX* GFP fluorescence distributions in microfluidic experiments
- Table S2: Fitted values for the 12 parameters of the model and statistical measurement of their robustness from bootstrap process
- Table S3: RT-qPCR primer sequences

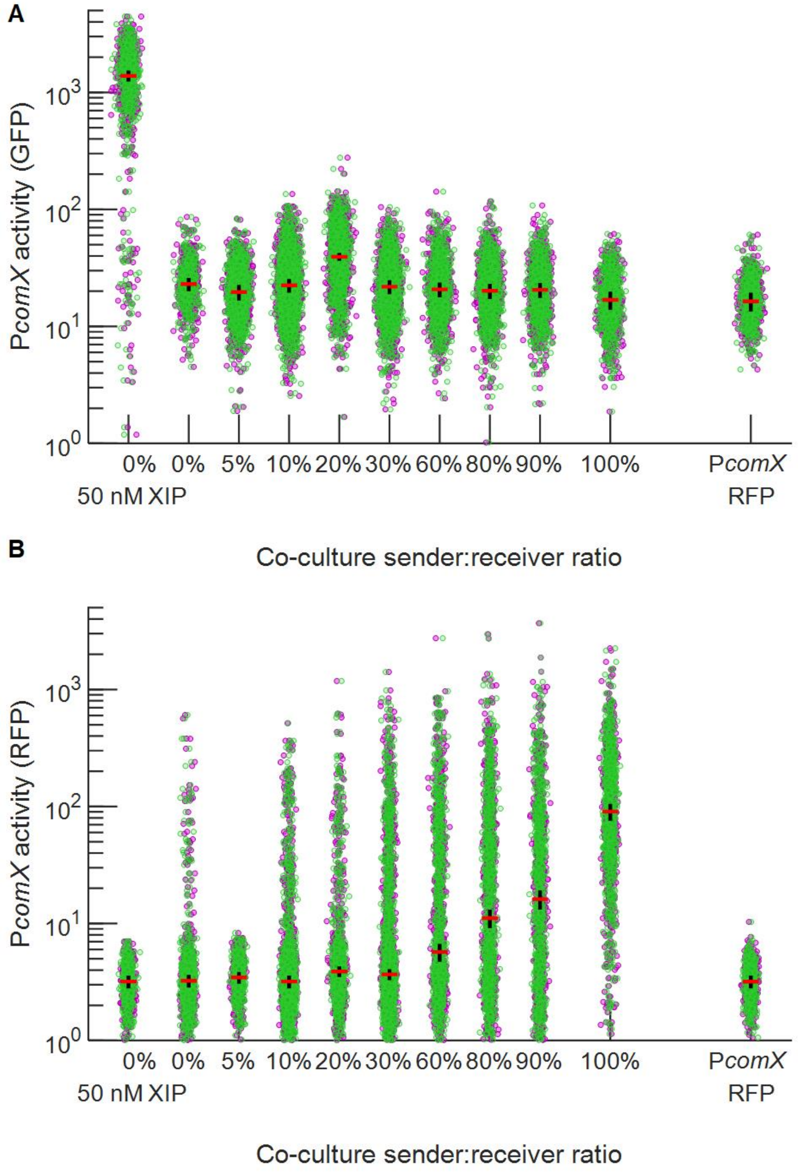


**Figure S1 – Response of cocultures is time-independent**

(A) GFP (*comX* reporter) and (B) RFP (*comY*) fluorescence of individual cells in co-cultures of sender (184comS P*comX-rfp*) and receiver (P*comX-gfp* Δ*comS*) strains of *S. mutans*. Samples are labeled by percentage by volume of 184*comS* (sender) culture in the initial preparation of the coculture. Fluorescence was measured immediately (0 h, green) after mixing, or 4 h (magenta) after mixing the coculture. The red horizontal bars show the median fluorescence immediately after mixing (0 h); the black vertical bars show the median at 4 h. Data are from the coculture experiment of Figure 4.

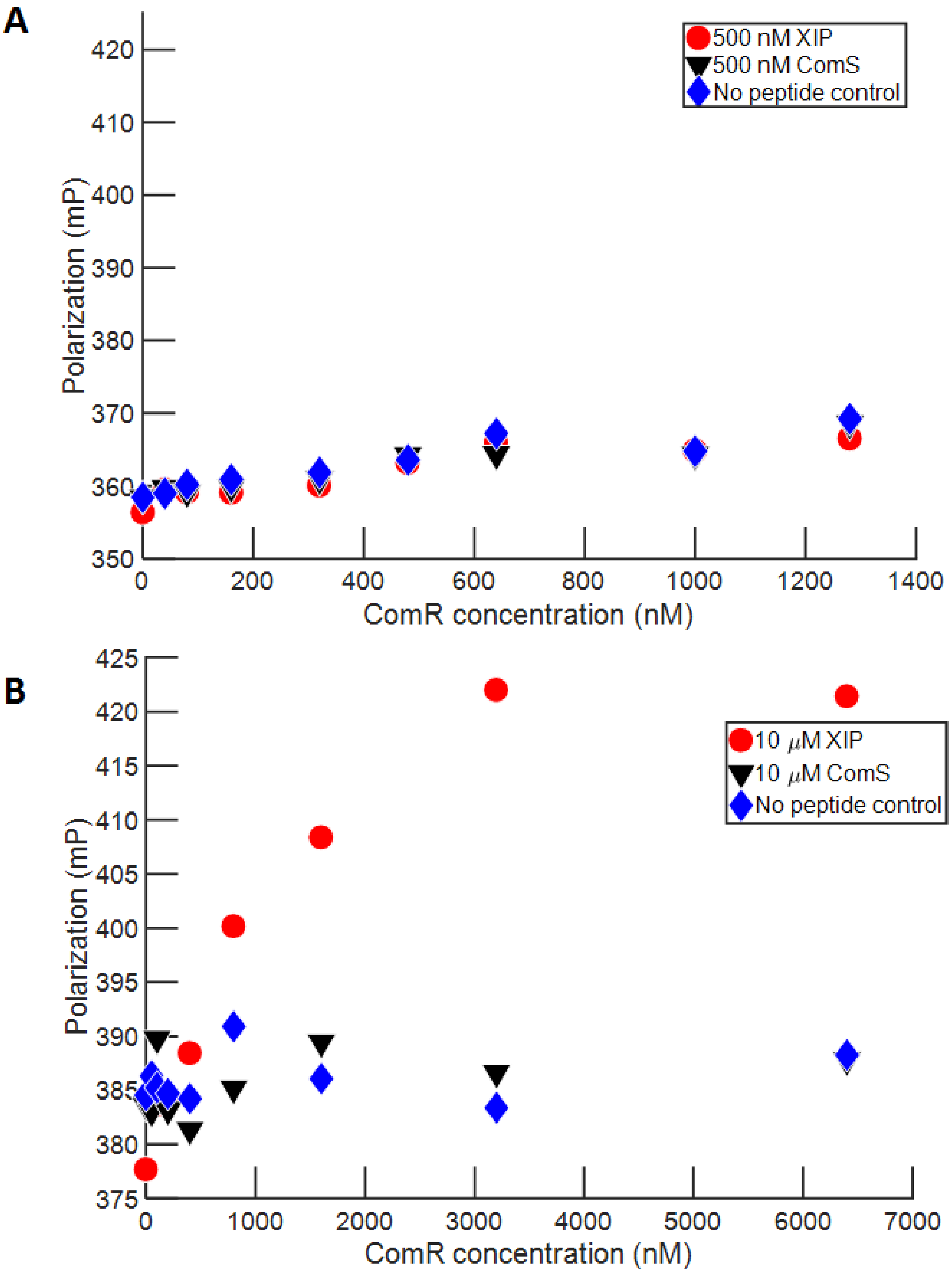


**Figure S2: Effect of histidine tag on ComR binding of ComS and XIP**

Fluorescence polarization study of ComS and XIP interaction with ComR that was N- or C-terminally tagged with 6X-histidine. Polarization is plotted versus [ComR] for (A) C-terminally tagged ComR and (B) N-terminally tagged ComR. In each case 1 nM fluorescent DNA and 0.05 mg ml-1 salmon DNA were present along with no signal peptides (black), XIP (blue) or ComS (green).

**Deterministic fit: Equation system and calculated parameter values**

The system of ODEs used to fit microfluidic data is given below. X represents ComX, Z the internal XIP concentration, S the internal ComS concentration, R the (constant) ComR concentration and Exo the exogenous XIP level. All units are in nM and seconds where appropriate. Other symbols are parameters describing the reaction kinetics. A star indicates one of the *V* parameters contributing to feedback, while unstarred *V*s indicate a maximum rate of production of ComX. Hill kinetics corresponding to inferred cooperativity from FP assays were used. Calculated parameters are given in Table S2. A 200-iteration bootstrap analysis of the data was performed in order to estimate parameter robustness, with the 10th and 90th percentiles of parameter values reported. These percentile values demonstrate preservation of the relative order of magnitude between dissociation constants for XIP-ComR and ComS-ComR.

$$\frac{dX}{dt} = \frac{V_1 R^2 Z^2}{R^2 Z^2 + K_X^4} + \frac{V_2 RS}{RS + K_S^2} - \beta_X X \quad (1)$$

$$\frac{dS}{dt} = \alpha_0 - \beta_S S - \gamma S + \frac{V_1^* R^2 Z^2}{R^2 Z^2 + K_X^4} + \frac{V_2^* RS}{RS + K_S^2} - \frac{V_2 RS}{RS + K_S^2} \quad (2)$$

$$\frac{dZ}{dt} = J(Exo - Z) - \beta_Z Z + \gamma S - \frac{V_1 R^2 Z^2}{R^2 Z^2 + K_X^4} - \frac{V_1^* R^2 Z^2}{R^2 Z^2 + K_X^4} \quad (3)$$

**Table S1: Parameters for gamma distribution fits to single cell P*comX* GFP fluorescence distributions in microfluidic experiments**.

| [XIP] | *a* | *b* |
|---|---|---|
| **Wild type** | - | - |
| 0 | 4.09 | 5.51 |
| 280 | 5.50 | 172 |
| 700 | 9.62 | 153 |
| 1840 | 11.0 | 184 |
| 3250 | 12.5 | 168 |
| 5230 | 12.4 | 174 |
| 6000 | 11.7 | 172 |
| ***ΔcomS*** | - | - |
| 0 | 6.14 | 2.29 |
| 30 | 7.10 | 2.44 |
| 850 | 4.24 | 166 |
| 940 | 5.11 | 148 |
| 3000 | 7.44 | 160 |
| 4020 | 7.61 | 172 |
| 6000 | 7.45 | 180 |

**Table S2: Fitted values for the 12 parameters of the model and statistical measurement of their robustness from bootstrap process.**

| Parameter | Best fit value – used for Fig. 7 | $10^{th}$ percentile from bootstrap | $90^{th}$ percentile from bootstrap | Units |
|---|---|---|---|---|
| $\alpha_0$ | 7.85 | 1.78 | 19.1 | nM $s^{-1}$ |
| $\beta_S$ | 7.17 x $10^{-3}$ | 7.97 x $10^{-4}$ | 1.32 x $10^{-2}$ | $s^{-1}$ |
| $\gamma$ | 0.452 | 0.227 | 2.74 | $s^{-1}$ |
| $V_1^*$ | 1.07 x $10^4$ | 3.8 x $10^3$ | 2.46 x $10^4$ | nM $s^{-1}$ |
| $V_2^*$ | 1.33 x $10^4$ | 3.22 x $10^3$ | 2.65 x $10^4$ | nM $s^{-1}$ |
| $K_x$ | 148 | 42 | 210 | nM |
| $K_S$ | 2740 | 849 | 3000 | nM |
| $V_1$ | 3.55 | 0.988 | 558 | nM $s^{-1}$ |
| $V_2$ | 778 | 309 | 2190 | $s^{-1}$ |
| $J$ | 9.39 | 6.14 x $10^{-2}$ | 17.6 | $s^{-1}$ |
| $\beta_Z$ | 1.28 | 0.264 | 3.07 | nM $s^{-1}$ |
| $\beta_X$ | 10.5 | 1.71 | 16.2 | $s^{-1}$ |

**Table S3: RT-qPCR primer sequences**

| Gene and direction | Primer sequence |
|---|---|
| *comX* forward | 5’-CGTCAGCAAGAAAGTCAGAAA C-3’ |
| *comX* reverse | 5’-ATACCGCCACTTGACAAACAG-3’ |
| *comS* forward | 5’-TCAAAAAGAAAGGAGAATAACA-3’ |
| *comS* reverse | 5’-TCATCTGAGATAAGGGCTGT-3’ |
| 16S rRNA forward | 5’-CACACCGCCCGTCACACC-3’ |
| 16S rRNA reverse | 5’-CAGCCGCACCTTCCGATACG-3’ |
